# A *mdg4* Retrotransposon Screen for X-linked Female Sterile Alleles and its Relationship with the Transcription Factor OVO

**DOI:** 10.64898/2026.02.12.705638

**Authors:** Leif Benner, Brian Oliver

## Abstract

In the germline, the *mdg4* retrotransposon integrates in close proximity to the location of OVO DNA binding motifs, suggesting that insertion bias is driven by the OVO transcription factor. A classical genetic example of this is the reversion of the dominant female-sterile allele, *ovo*^*D1*^, by the transposition of *mdg4* into the *ovo* promoter where OVO protein binds. We wanted to take advantage of this relationship and determine if we could recover female sterile alleles along the X chromosome due to *mdg4* insertion, with the hypothesis that these would be genes that OVO binds and transcriptionally regulates in the germline. We mobilized the *mdg4* retrotransposon with the use of mutants for the lncRNA gene *flamenco* (*flam)* and recovered 17 recessive female sterile alleles out of a total of 1,192 chromosomes screened. We identified 11 complementation groups, for which a *mdg4* insertion was responsible for female sterility in 7 groups. Notably, a complementation group consisting of 6 alleles was found to be the result of a *Doc* transposable element insertion into the gene *Grip91* and is potentially evidence for a *Doc* insertional ‘hotspot’ in the genome. Our screen also uncovered that 7/17 recessive female sterile chromosomes contained multiple transposable element insertions indicating that *flam*^−^ females derepress numerous transposable elements that can lead to multiple transposon insertions along a single chromosome, as has been suggested previously. Altogether, we found that *mdg4* did have an insertion bias into OVO bound regions of the genome that can result in female sterility, however, this was the case for a minority of the female sterile alleles recovered with this method.

**Article Summary:** The retrotransposon *mdg4* preferentially inserts near binding sites of the female germline transcription factor OVO in *Drosophila melanogaster*, most notably at the *ovo* locus itself. We leveraged this relationship to screen for X-linked recessive female-sterile mutations generated by *mdg4* mobilization in *flamenco* mutant females. From 1,192 chromosomes, we recovered 17 female-sterile alleles across 11 complementation groups. *mdg4* insertions were significantly enriched in OVO-bound regions but accounted for only a subset of sterility phenotypes, revealing substantial background mutagenesis by other transposable elements. These results refine the OVO-*mdg4* relationship and highlight both the promise and limitations of transposon-based genetic screens.

## Introduction

The *mdg4* retrotransposon (a.k.a. *gypsy*) shares a long scientific history with the transcription factor OVO. The *ovo* locus was first identified as a set of three dominant female sterile alleles along the X chromosome (*ovo*^*D1*^, *ovo*^*D2*^, *ovo*^*D3*^)(Busson et al., 1983; Komitopoulou et al., 1983). In a serendipitous fashion, it was found that when *ovo*^*D1*^ males were crossed to a ‘wild type’ *y*^*1*^ *v*^*1*^ *f*^*1*^ *mal*^*1*^ stock, transheterozygous female progeny had a high rate of *ovo*^*D1*^ reversion from dominance to loss-of-function, therefore, heterozygous females were fertile and *ovo*^*D1*^ had lost its antimorphic behavior (Busson et al., 1983). When other female genotypes were crossed to *ovo*^*D1*^ males, there was no detectable reversion of *ovo*^*D1*^ in the progeny, indicating that there was a unique maternal factor in *y*^*1*^ *v*^*1*^ *f*^*1*^ *mal*^*1*^ females causing *ovo*^*D1*^ to revert. Future work determined that reversion of *ovo*^*D1*^ was due to the insertion of *mdg4* and *copia* transposable elements into *ovo*^*D1*^, and that these elements had mobilized in *y*^*1*^ *v*^*1*^ *f*^*1*^ *mal*^*1*^ females (Mével-Ninio et al., 1989). It was also determined that the mobilization of *mdg4* was genetically linked to a region along the *y*^*1*^ *v*^*1*^ *f*^*1*^ *mal*^*1*^ X-chromosome, indicating that there was likely a locus responsible for silencing the transposition of *mdg4*. This indeed was found to be the case and the resulting locus was called *flamenco* (*flam*) (Pélisson et al., 1994; Prud’homme et al., 1995). *flam* was later found to encode a RNA transcript that is processed into piRNAs and is the mechanism through which *flam* silenced the activity of transposons such as *mdg4* (Brennecke et al., 2007; Sarot et al., 2004)(a history of *ovo, flamenco, mdg4*, and the birth of the piRNA field is nicely described in Coline et al., 2014).

It turned out that *mdg4* integration into *ovo*^*D1*^ was not random, but that the *ovo* DNA region conferred a specificity for *mdg4* integration. OVO protein binds to the *ovo* DNA region proximal to the promoter (Andrews et al., 2000; Benner et al., 2024; Bielinska et al., 2005; Lü et al., 1998) and most of the *mdg4* insertions into *ovo*^*D1*^ were clustered in this region (Dej et al., 1998; Labrador & Corces, 2001; Mével-Ninio et al., 1989; Song et al., 1994). An elegant set of experiments showed that *mdg4* had a preference for inserting near OVO DNA binding motifs by creating a reporter with an array of high affinity OVO DNA binding motifs, nested between enhancers and the promoter for the *yellow* gene (Labrador et al., 2008). It was found that *mdg4* inserted frequently, and in close proximity, to the array of OVO motifs. Mutating these motifs to a sequence that OVO did not bind abolished the ability for *mdg4* to insert into this reporter. Clearly, *mdg4* insertions were biased, and likely driven by where OVO was binding in the genome, either by directly associating with OVO or taking advantage of open chromatin in the germline due to OVO binding.

OVO is a required transcription factor in the female germline and has been shown to regulate the expression of hundreds of germ cell-specific genes (Benner et al., 2024; Oliver et al., 1987, 1990). Understanding not only the repertoire of putative OVO target genes required in the female germline, but also how OVO regulates target genes, is an important area of research in germ cell biology. In order to further characterize the transcriptional program downstream of OVO, we were interested in taking advantage of the relationship between OVO binding and *mdg4* insertional bias by conducting a screen for recessive female sterility due to *mdg4*. Presumably, if *mdg4* inserts proximal to genes that are bound by OVO and are required for female fertility, then we could recover genes that are transcriptionally regulated by OVO and are functionally required in the female germline. If this relationship is true, then this would be a reliable and unbiased method to confirm previously identified putative OVO target genes.

Although there is positive evidence for a relationship between OVO DNA binding and *mdg4* insertion, there are potential drawbacks to conducting a screen in this manner. For example, mobilizing *mdg4* elements is dependent on using *flam*^−^ females. *flam* is responsible for the regulation of other transposable elements in addition to *mdg4*, and using *flam* mutants might have the unintended consequence of recovering female sterile alleles due to the insertion of transposable elements other than *mdg4* (Desset et al., 2003; Mével-Ninio et al., 1989, 2007; Song et al., 1994; Zanni et al., 2013). Also, *mdg4* element insertions are sometimes genomically unstable (Mizrokhi et al., 1985; Prud’homme et al., 1995). It is possible to isolate female sterile alleles due to *mdg4* that would eventually revert and might be difficult to identify if reversion occurs prior to DNA-sequencing. Regardless, we decided to test this directly and screen for *mdg4* induced recessive female sterile alleles along the X-chromosome and assess if this would be an efficient screening method to employ at a larger scale. We also wanted to further determine if there is a relationship between *mdg4* insertions and OVO DNA binding by comparing the location of *mdg4* insertions to recently published OVO ChIP-seq data.

## Methods

### Fly Husbandry

All stocks used in this study are listed in Flybase Author Reagent Table (Table S1). Flies were maintained on glucose fly media (Archon Scientific, Durham, NC) and crosses were completed at 25°C with 65% relative humidity and constant light. All alleles used in this study can be found in the Flybase ART Table (Table S1).

### Screen

Before beginning our *mdg4* female sterile screen, we wanted to confirm active *mdg4* mobilization in transheterozygous combinations of *flam*^−^ alleles. A classical method to test for *mdg4* mobilization is by taking advantage of the *mdg4* retrotransposons propensity to insert into the promoter region of the dominant-female sterile *ovo*^*D1*^ allele (Mével-Ninio et al., 1989). *mdg4* insertion into *ovo*^*D1*^ reverts the dominant-female sterility of this allele and results in fertile females. We performed this assay by crossing *ovo*^*D1*^ males to two different combinations of *flam*^−^ females (Figure S1). We were able to recover fertile *ovo*^*D1*^ transheterozygous revertants from both *flam*^−^ combinations while we did not recover *ovo*^*D1*^ revertants in the *flam*^−^/+ control female crosses. We performed whole genome DNA sequencing on two revertant males and confirmed *mdg4* element insertion into the first intron of *ovo*^*D1*^ (*ovo-RA* transcript).

The cross scheme to mobilize the *mdg4* retrotransposon and isolate female sterile alleles along the X chromosome are outlined in Figure 1A. *mdg4* mobilization occurs in *y*^*1*^ *v*^*1*^ *f*^*1*^ *mal*^*1*^ *lncRNA:flam*^*1*^ *su(f)*^*1*^/*FM7c, sn*^*+*^ *P*{*lyB*}*lncRNA:flam*^*py+P*^ adult females of cross G0 and *mdg4* retrovirus particles are maternally inherited by the offspring of cross G1 (Kim et al., 1994; Pélisson et al., 1994; Song et al., 1994, 1997). *flam*^−^ females from cross G0 were only mated up to 4 days post-eclosion to maximize the number of potential *mdg4* insertions as has been previously reported (Prud’homme et al., 1995). *mdg4* integration events can occur in the germline along the *y*^*1*^ *cv*^*1*^ *v*^*1*^ *f*^*1*^ chromosome (denoted by *fs(1)*^*\**^) in these females. Also notable, in cross G2, females with the genotypes *C(1)DX, y*^*1*^ *f*^*1*^*/Y* and *FM7c, cv*^*WJB*^ *sn*^*+*^ *f*^*PDB*^*/w*^*1118*^ *sov*^*ML150*^ were crossed to a single *fs(1)*^*\**^, *y*^*1*^ *cv*^*1*^ *v*^*1*^ *f*^*1*^ male within the same vial. The genotypes of all the resulting progeny are easily distinguishable from each other based on their respective recessively linked mutations and allowed us to skip a generation for the fertility tests.

**Figure 1:**
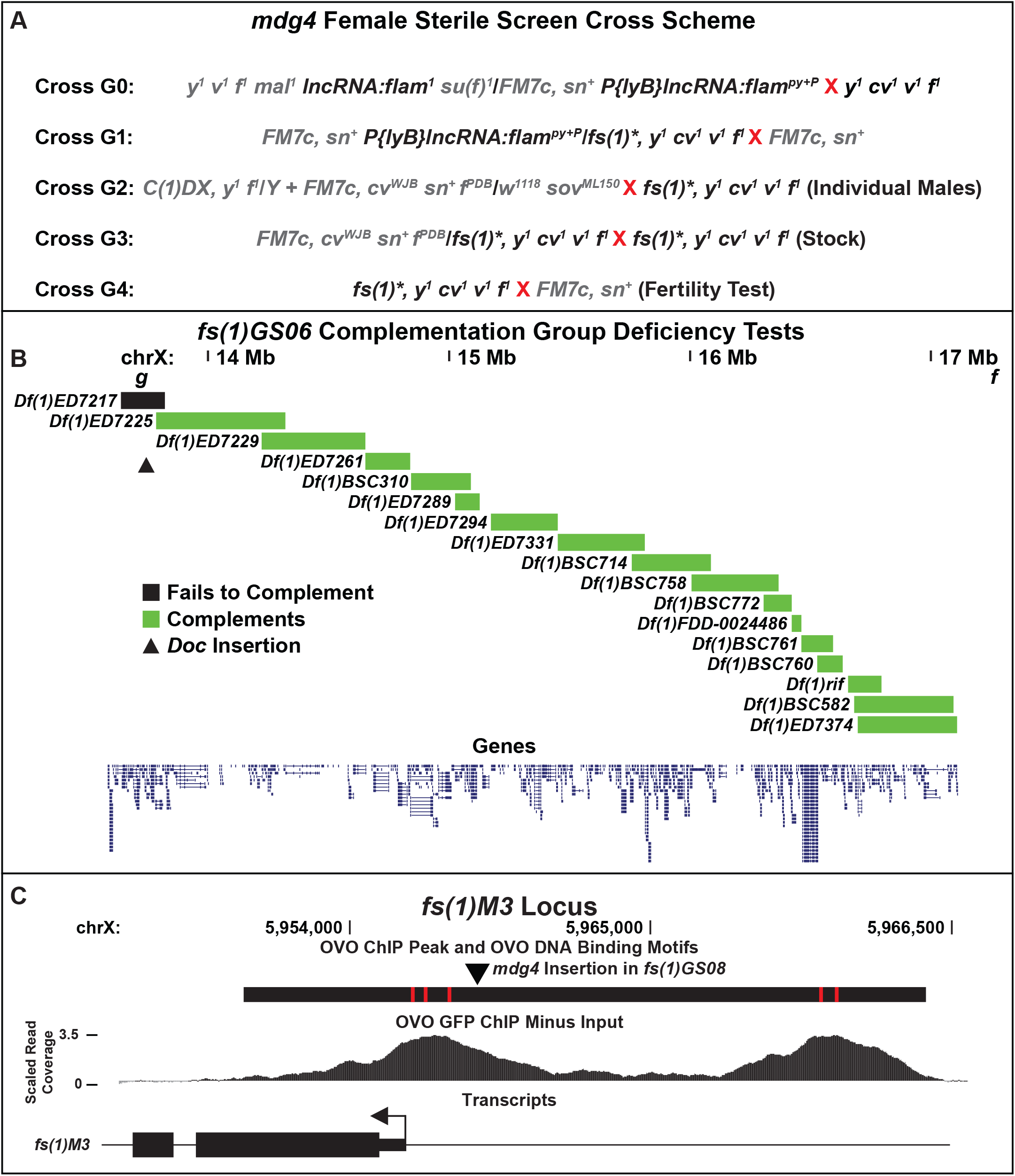
Cross scheme, deficiency testing, and *mdg4* insertion in *fs(1)M3*. A) Cross scheme to recover recessive female sterile alleles from *flam*^−^ females. B) Deficiency mapping of *fs(1)GS06*. Boxes indicate the deletion interval for each deficiency. Green boxes indicate that the deficiency complemented *fs(1)GS06*, black box indicates the deficiency that failed to complement *fs(1)GS06*, and the triangle indicates the location of the *Doc* insertion into *Grip91*. C) Genomic region of the *mdg4* insertion upstream of *fs(1)M3*. Large black box indicates the significant OVO ChIP peak from the below OVO GFP ChIP minus input scaled read coverage. Red boxes indicate significant OVO DNA binding motifs and the triangle indicates the location of the *mdg4* insertion. The *fs(1)M3* transcript is indicated below with the arrow signifying the TSS.

### Meiotic Mapping and Complementation Tests

*fs(1)*^*\**^, *y*^*1*^ *cv*^*1*^ *v*^*1*^ *f*^*1*^ males were crossed to *OregonR* (wildtype) females to allow for recombination along the X chromosome. A total of 5 individual *fs(1)*^*\**^ recombinant males of the following genotypic intervals, *y*^*1*^, *y*^*1*^ *cv*^*1*^, *y*^*1*^ *cv*^*1*^ *v*^*1*^, *cv*^*1*^ *v*^*1*^ *f*^*1*^, *v*^*1*^ *f*^*1*^, *f*^*1*^, were crossed back to the parental *fs(1)*^*\**^, *y*^*1*^ *cv*^*1*^ *v*^*1*^ *f*^*1*^/*FM7c, cv*^*WJB*^ *sn*^*+*^ *f*^*PDB*^ females. A single vial consisting of 10 transheterozygous *fs(1)*^*\**^ recombinant/*fs(1)*^*\**^, *y*^*1*^ *cv*^*1*^ *v*^*1*^ *f*^*1*^ females were tested for fertility by crossing to *FM7c, sn*^*+*^ males. The same fertility tests were carried out with *fs(1)*^*\**^, *y*^*1*^ *cv*^*1*^ *v*^*1*^ *f*^*1*^/*Df(1)* females.

### DNA-sequencing and Transposable Element Insertion Analysis

For each female sterile and lethal allele, genomic DNA was isolated from 30 females with the Qiagen blood and tissue kit (Hilden, Germany) utilizing the manufacturers insect protocol. Library preparation and sequencing was carried out as described before (Hammond et al., 2020). *fs(1)GS03* was initially a failed library and was re-prepped and sequenced with different methods as described previously (Muron et al., 2022). Paired-end reads were mapped with Hisat2 (Kim et al., 2019) to a masked version of the *Drosophila melanogaster* dm6 release (Hoskins et al., 2015; Smit et al., 2015) supplemented with the DNA sequences of all published transposable elements (Kaminker et al., 2002). Mapped reads were sorted and indexed with Samtools (Danecek et al., 2021). DNA-seq data of all alleles generated in this work are available at SRA:PRJNA1422399.

In order to identify the locations of the transposable elements, we extracted all discordant read pairs mapping to the X chromosome and any transposable element. For each X chromosome discordant read, we created a 1000 NT genomic interval based on each read’s genomic location with bedtools slop (Quinlan & Hall, 2010) and then collapsed all overlapping intervals for a given transposable element with bedtools merge, giving us a unique genomic interval where a possible transposable element existed on the X chromosome. For each female sterile allele, unique transposable element locations were intersected with the control chromosome with bedtools intersect and transposable element locations that existed along both the female sterile and control chromosome were removed. This left us with unique transposable element locations for each female sterile allele. All potential unique insertions were manually checked for accuracy in IGV (Robinson et al., 2011). We further defined a transposable element insertion with the following characteristics. There had to be evidence of discordant read pairs on both sides of the insertion site and there also had to be an evident loss of read coverage at the insertion site indicating a transposition of novel DNA sequences. Insertions that did not meet these criteria were not examined further.

Figure 1B and 1C utilized the UCSC genome browser for visualization (Casper et al., 2026).

### *mdg4* and OVO Enrichment Analysis

Significant OVO ChIP peaks were determined as previously described (Benner et al., 2024) utilizing the ‘OVO-GFP’ ChIP-seq datasets only. ATAC-seq datasets from germline stem cells and stage 5 nurse cells (DeLuca et al., 2020) were analyzed identically as before (Benner et al., 2024). Bigwig files of OVO ChIP-seq datasets were generated with deeptools software (Ramírez et al., 2016). For enrichment analysis, coordinates from significant OVO ChIP peaks, germline stem cell, and stage 5 nurse cell ATAC-seq peaks along the X chromosome were randomly shuffled 1000 individual times with bedtools shuffle (-noOverlapping). *mdg4* and *Doc* insertion locations were then intersected with the 1000 randomly shuffled peak locations with bedtools intersect. To calculate a p-value for enrichment, the sum of randomly shuffled sequences that contained an equal or greater number of intersections compared to the actual dataset was then divided by 1000.

## Results and Discussion

### Isolation and Meiotic Mapping of fs(1) Alleles

Mutants for the lncRNA gene *flamenco* (*flam*) have classically been used to allow for the mobilization of the *mdg4* retrotransposon (Dej et al., 1998; Kim et al., 1994; Labrador et al., 2008; Labrador & Corces, 2001; Mével-Ninio et al., 1989; Pélisson et al., 1994; Prud’homme et al., 1995; Song et al., 1994, 1997). This is because RNAs expressed from *flam* are processed into piRNAs that directly repress the expression of *mdg4* (Brennecke et al., 2007; Sarot et al., 2004). Therefore, mutations in *flam*, that result in the loss of these piRNAs, leads to a derepression of *mdg4* and subsequent mobilization. Interestingly, this regulation occurs in the somatic cells of the ovary and loss of *flam* leads to the production of *mdg4* retroviruses that invade the female germline and become maternally deposited (Kim et al., 1994; Pélisson et al., 1994; Song et al., 1994, 1997). This leads to the potential for *mdg4* insertion and mutagenesis in the offspring of these mothers.

Before beginning our screen, we wanted to confirm *mdg4* mobilization in *y*^*1*^ *v*^*1*^ *f*^*1*^ *mal*^*1*^ *lncRNA:flam*^*1*^ *su(f)*^*1*^/*FM7c, sn*^*+*^ *P*{*lyB*}*lncRNA:flam*^*py+P*^ females. In order to test this, we crossed *ovo*^*D1*^ males to *y*^*1*^ *v*^*1*^ *f*^*1*^ *mal*^*1*^ *lncRNA:flam*^*1*^ *su(f)*^*1*^/*FM7c, sn*^*+*^ *P*{*lyB*}*lncRNA:flam*^*py+P*^ females and scored the number of fertile *ovo*^*D1*^ revertant chromosomes (Figure S1). We had a 2.1% reversion rate of *ovo*^*D1*^, which we confirmed was due to *mdg4* through DNA-sequencing, compared to controls that did not have any revertant *ovo*^*D1*^ chromosomes. Therefore, there was sufficient *mdg4* mobilization in these transheterozygous *flam*^−^ females to conduct our screen.

For our screen, we decided to isolate female sterile mutations due to *mdg4* transposition along the X chromosome. Since previous work showed that *mdg4* transposes in close proximity to OVO DNA binding motifs (Labrador et al., 2008; Labrador & Corces, 2001), a *mdg4* insertion causing female sterility is potentially an OVO target gene. From our screen (Figure 1A), we isolated 1,192 individual X chromosomes with the recovery of 16 female-specific sterile allele and one female-specific lethal allele. Therefore, we had a female-specific sterile recovery rate of 1.42%. We also had 89 F2 males that were sterile and did not score male-specific lethality in this screen.

Next, we meiotically mapped the *fs(1)* loci by crossing *fs(1), y*^*1*^ *cv*^*1*^ *v*^*1*^ *f*^*1*^ males to wild type females (*oregonR*) and selected for the following recombinants in the next generation: *y*^*1*^, *y*^*1*^ *cv*^*1*^, *y*^*1*^ *cv*^*1*^ *v*^*1*^, *cv*^*1*^ *v*^*1*^ *f*^*1*^, *v*^*1*^ *f*^*1*^, *f*^*1*^. Five individual males accounting for each interval were crossed back to the original *fs(1), y*^*1*^ *cv*^*1*^ *v*^*1*^ *f*^*1*^ parental chromosome and tested for sterility. This allowed us to roughly map the location of the *fs(1)* locus for our eventual sequencing analysis (Table S2). We were not able to confidently meiotically map *fs(1)GS05* and *fs(1)GS15* due to inconclusive sterility results of different recombinants. Although we were not sure initially as to the reason behind this result, this was our first indication that we potentially had multiple mutations along the chromosome, which we will discuss below.

We next complementation tested all *fs(1)* alleles that were meiotically mapped within a specific region along the X chromosome. For example, all *fs(1)* alleles that were mapped between *y* and *cv* were complementation tested. The two *fs(1)* alleles that had inconclusive mapping results were complementation tested to all *fs(1)* alleles. Complementation results indicated that we had 11 unique complementation groups represented by a total of 17 alleles. With *fs(1)GS06, 07, 09, 14, 15, 16, 17* (we will refer to this complementation group as *fs(1)GS06*) all failing to complement each other and therefore were likely a single complementation group for the same female sterile locus. Complementation results can be found in Table S2.

### DNA-seq of fs(1) Alleles

In order to determine the mutations resulting in female sterility, we conducted 50bp paired-end sequencing on all *fs(1)* alleles recovered from the screen as well as the original non-mutagenized *y*^*1*^ *cv*^*1*^ *v*^*1*^ *f*^*1*^ chromosome. This gave us the ability to determine novel transposable element insertions compared to our background non-mutagenized genotype. We elected to do whole genome sequencing instead of a more directed approach, such as inverse-PCR sequencing with primers targeting *mdg4*, since we were unsure if all the female sterile mutations would be a direct result of *mdg4* transposition and not another transposable element. Thus, we could discern the potential mobilization of non-*mdg4* elements in *flam*^−^ mutants.

Our sequencing results confirmed that 8/11 complementation groups contained a novel *mdg4* transposable element insertion and 7/11 were within the genomic window consistent with our meiotic mapping results for sterility (Table 1). Notably, we could not identify a *mdg4* insertion in *fs(1)GS03, fs(1)GS10*, and the complementation group *fs(1)GS06* (more below). Relevant information for *fs(1)* alleles due to *mdg4* insertions are listed in Table 1. Information about *fs(1)* alleles with non-*mdg4* insertions are listed in Table S3.

**Table 1:**
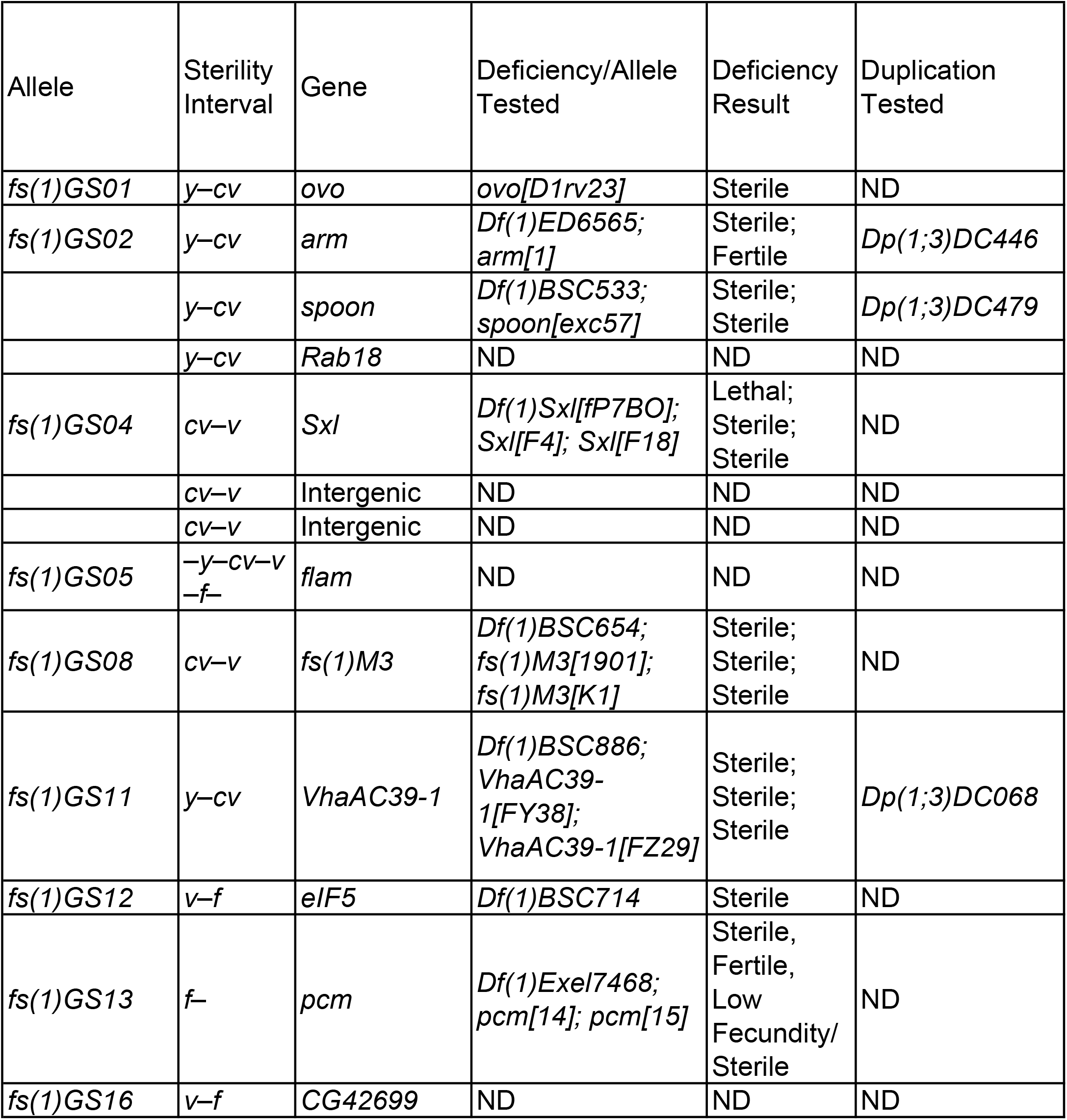
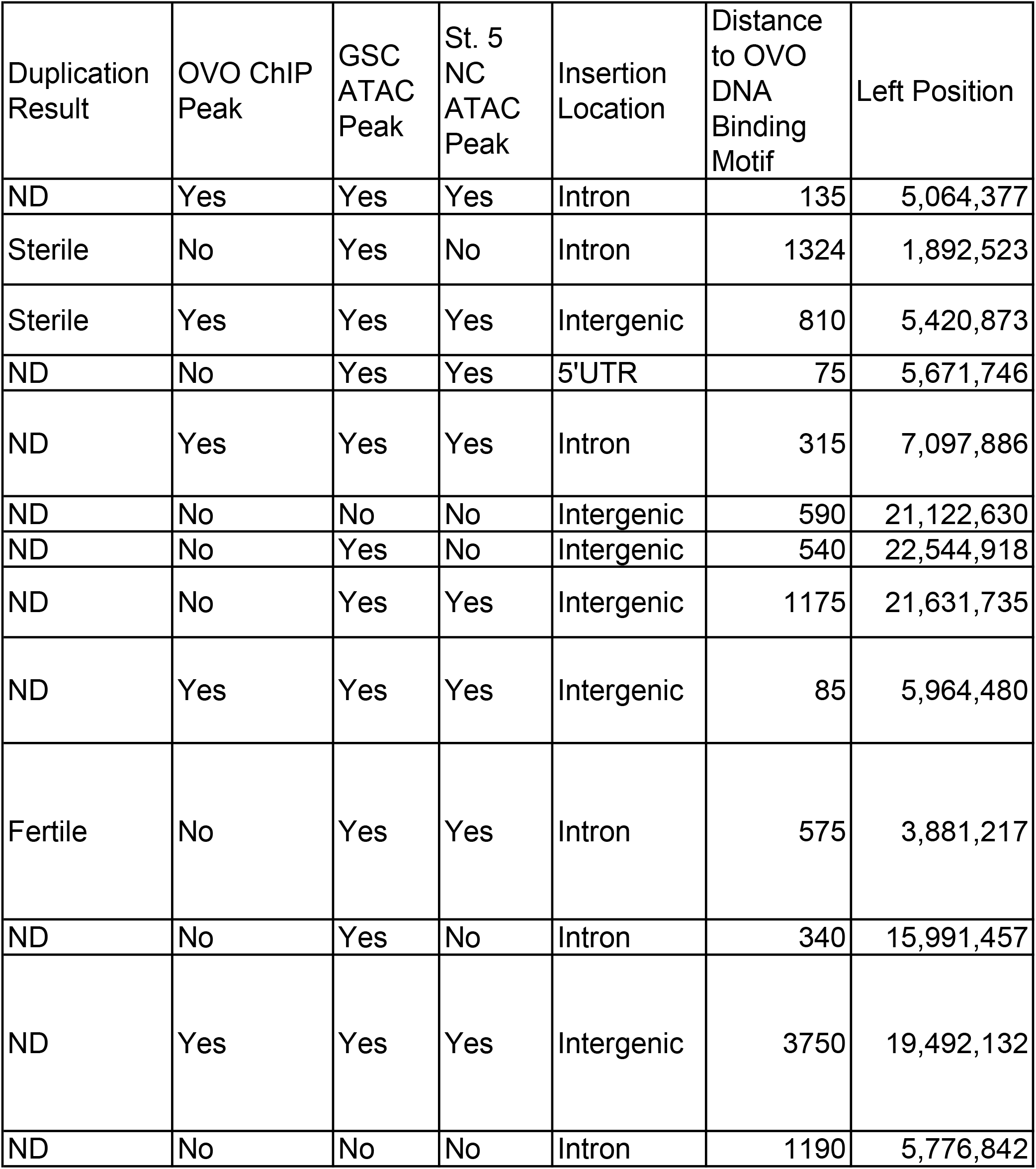
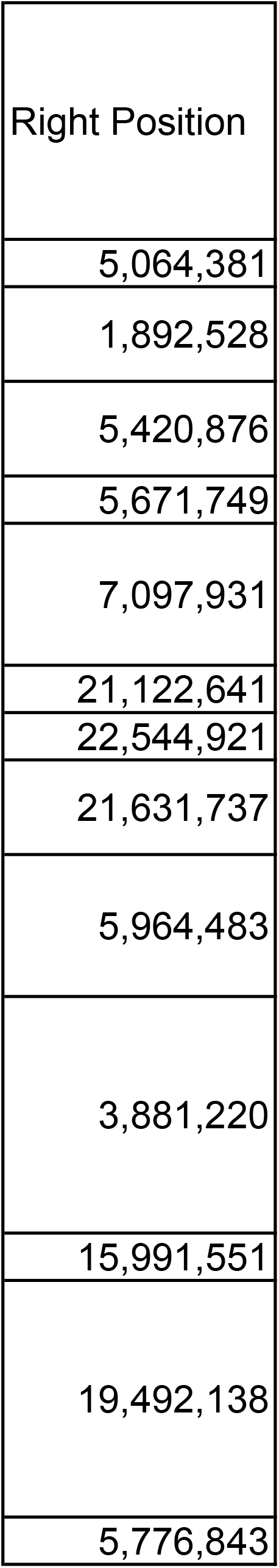
Characteristics of all *mdg4* dependent recessive female sterile alleles.

### Description of *fs(1)* Alleles

#### fs(1)GS01

*fs(1)GS01* sterility mapped to the *y*-*cv* interval and we found a novel *mdg4* insertion within this interval between the first and second exon of *ovo-B* (based on the *ovo-RA* transcript (Öztürk-Çolak et al., 2024)). This insertion was within 40 nucleotides of the two *mdg4* insertions found in the *ovo*^*D1*^ revertants recovered earlier. This region is the previously described ‘hotspot’ for *mdg4* insertions that revert *ovo*^*D1*^ to fertility (Dej et al., 1998; Labrador & Corces, 2001; Mével-Ninio et al., 1989; Song et al., 1994). *fs(1)GS01* failed to complement *ovo*^*D1rv23*^ sterility and therefore this *mdg4* insertion indeed disrupted *ovo* gene activity. The *mdg4* insertion was also found to be within a significant OVO ChIP peak, as well as an open chromatin region in both germline stem cell (GSC) ATAC-seq and stage 5 nurse cell (St. 5 NC) ATAC-seq data. The closest OVO DNA binding motif to this insertion site was roughly 135 NTs upstream.

We also had evidence for a novel *Doc* insertion within the 3’UTR of the gene *Imp* along this chromosome. However, this insertion is outside of the *y*-*cv* interval responsible for sterility and therefore this *Doc* insertion was unlikely to contribute to the sterility phenotype.

#### fs(1)GS02

*fs(1)GS02* sterility mapped to the *y*-*cv* interval and there was evidence for three novel *mdg4* insertions within this region. One *mdg4* insertion was into the last intron of *armadillo* (*arm*), the second insertion was slightly upstream of the gene *spoonbill* (*spoon*), and the third *mdg4* insertion was in the 5’UTR of *Rab18*. Mutants for both *arm* and *spoon* have previously been found to result in female sterility and in both cases, sterility is germline-dependent (Neuman-Silberberg, 2007; Peifer et al., 1993; Perrimon et al., 1989a; Perrimon & Mahowald, 1987; White et al., 1998). *fs(1)GS02* failed to complement a deficiency including *arm*, (*Df(1)ED6565*), however, *fs(1)GS02* did complement lethality and fertility of *arm*^*1*^, indicating that the insertion might not be disrupting the activity of *arm*, and possibly disrupting the activity of a nearby gene that is required for female fertility. A deficiency including *spoon* (*Df(1)BSC533*), and female sterile allele *spoon*^*exc57*^, also failed to complement *fs(1)GS02* sterility suggesting that this insertion is disrupting the activity of *spoon* and leading to female sterility. Duplications for both *arm* (*Dp(1;3)DC446*) and *spoon* (*Dp(1;3)DC479*) independently failed to rescue *fs(1)GS02* homozygous sterility, consistent with the complementation results that indicate each insertion is independently resulting in female sterility. We could not test if the *mdg4* insertion within *Rab18* also leads to sterility since the gene downstream of *Rab18, RpL35*, is a haploinsufficient locus and therefore there are no deficiencies for this region.

The *mdg4* insertion in *arm* overlapped a GSC ATAC peak but did not overlap an OVO ChIP or St. 5 NC ATAC peak. However, OVO ChIP data indicates that OVO does bind to *arm* at two discrete locations. One peak overlaps the *arm* promoter and another peak overlaps the second to last intron/exon. The OVO ChIP peak maxima/DNA binding motif overlapping the second to last intron/exon is roughly 1,324 NTs away from the *mdg4* insertion site. The *mdg4* insertion within *spoon* was 8 NTs upstream of the *spoon-RA* transcriptional start site (TSS). This insertion overlapped an OVO ChIP, GSC ATAC, and St. 5 NC ATAC peak and was 810 NTs away from a significant OVO DNA binding motif. Although we could not definitively test whether the *mdg4* insertion in *Rab18* also resulted in sterility, the *mdg4* insertion overlapped a GSC and St. 5 ATAC peak, but not an OVO ChIP peak. There are no OVO ChIP peaks overlapping *Rab18*, but the *mdg4* insertion is 75 NTs away from an OVO DNA binding motif.

#### fs(1)GS03

*fs(1)GS03* sterility mapped to the *v*-*f*-Rt interval and we did not have any evidence within our sequencing data for a novel *mdg4* insertion along this chromosome. Within this interval we were able to detect a novel *hobo* element insertion within the 3’UTR of the gene *Karl* and a *17*.*6* element insertion into the first intron of *U2af50. fs(1)GS03* complemented a deficiency including *Karl* (*Df(1)BSC541*) but failed to complement a deficiency including *U2af50* (*Df(1)FDD-0089495*), indicating that the insertion within *U2af50* was likely leading to female sterility. Knockdown of *U2af50* in the female germline has been previously shown to result in female sterility (Yan & Perrimon, 2015).

#### fl(1)GS04

*fl(1)GS04* was our only female lethal allele recovered from our screen and lethality mapped to the *cv*-*v* interval. Within this interval we detected a novel *mdg4* insertion in the first intron of the gene *Sex-lethal* (*Sxl*, location according to the *Sxl-RD* transcript). This insertion is downstream of the *Sxl* ‘maintenance’ promoter, which is the promoter utilized by *Sxl* transcripts which are eventually involved in the female-specific splicing of *transformer* and *male-specific lethal 2* genes in somatic tissues (Salz & Erickson, 2010). The female lethal deficiency *Sxl*^*fP7B0*^ (Salz et al., 1987) failed to complement lethality of *fl(1)GS04*. Furthermore, the female viable, but germline-dependent sterile alleles *Sxl*^*F4*^ and *Sxl*^*F18*^ (Mohler & Carroll, 1984; Oliver et al., 1990) complemented lethality but failed to complement sterility of *fl(1)GS04*. Altogether indicating that the insertion was disrupting *Sxl* activity in both somatic and germline cells. This *mdg4* insertion was found within an OVO ChIP, GSC ATAC, and St. 5 NC ATAC peak and was 315 NTs away from an OVO DNA binding motif. Interestingly, *Sxl* is also genetically downstream of OVO in the female germline sex determination pathway and this insertion raises the possibility that OVO might directly regulate *Sxl* expression in the female germline (Oliver et al., 1990, 1993; Oliver & Pauli, 1998).

We identified two additional intergenic *mdg4* insertions along this chromosome to the right of *f*, however, since lethality mapped to the *cv*-*v* region we do not believe these insertions contribute to the lethality phenotype. There were also two *Doc* insertions to the left of *cv* along this chromosome. One insertion was within the introns of overlapping genes *rg* and *CG15465*, another, within the gene *lncRNA:CR45519*. An additional intergenic *Tirant* insertion was detected to the right of *v*. For the same reason above, we do not believe any of these insertions are deleterious to viability.

#### fs(1)GS05

Meiotic mapping of *fs(1)GS05* sterility produced inconclusive results. All meiotic intervals tested were fertile. This possibly indicated that there were two mutations along this chromosome that genetically interacted resulting in synthetic sterility. However, they would have to be distally located. Regardless, our DNA-seq data only contained evidence for a single transposable element insertion, which was a *mdg4* insertion 160 NTs upstream of the annotated *flam* TSS. *flam* is to the right of *f* along the X chromosome and since this interval was not homozygous sterile based on our meiotic mapping, we did not test this allele against a *flam*^−^ chromosome or deficiency (since the potential other mutation would be heterozygous and therefore a negative complementation result is uninformative). Since this *mdg4* insertion in *flam* is proximal to the centromere of the acrocentric X, this might be evidence for our inability to recover sterile recombinants if there is indeed a second interacting mutation along this chromosome. However, we could not identify a second transposable element insertion along this chromosome and were therefore unable to resolve the cause of sterility.

#### fs(1)GS06 (also fs(1)GS07, fs(1)GS09, fs(1)GS14-17)

*fs(1)GS06* was part of the complementation group consisting of six other alleles. Meiotic mapping of *fs(1)GS06* placed the female sterility between *v* and *f*. This was also the case for all other members of this complementation group except *fs(1)GS15*, which produced inconclusive meiotic mapping results. Interestingly, we could not detect a transposable element insertion within this region in our DNA-seq data for any of these alleles. Perplexed by this, we refined the meiotic mapping to determine that sterility was to the right of *garnet* (*g*) and therefore between the *g* and *f* interval. We decided to perform complementation tests with deficiencies spanning this interval and found that one deficiency, *Df(1)ED7217*, failed to complement sterility of *fs(1)GS06* (Figure 1B). A manual search of the sequencing data within this deficiency region allowed us to identify a single *Doc* transposable element insertion within the first intron of *Grip91*, a known female sterile gene (Barbosa et al., 2000). This insertion also existed in all alleles of this complementation group. Furthermore, the *fs(1)GS06* failed to complement lethality of *Grip91*^*1*^ (Perrimon et al., 1989b) and was found to complement lethality of *Grip91*^*xr16*^ (Barbosa et al., 2000), however, *fs(1)GS06*/ *Grip91*^*xr16*^ females were sterile. Altogether, this is strong evidence that this *Doc* insertion is disrupting *Grip91* activity and causal for female sterility.

The reason we were not initially able to detect this insertion is that in the DNA-seq libraries from our non-mutagenized control *y*^*1*^ *cv*^*1*^ *v*^*1*^ *f*^*1*^ stock, there was also evidence of reads that mapped to the *Doc* transposable element at this location. However, although there were reads mapping to the *Doc* transposable element, there was still an equal read depth coverage across the potential insertion site in *y*^*1*^ *cv*^*1*^ *v*^*1*^ *f*^*1*^ DNA-seq libraries, which would indicate that there is not an insertion in the contiguous genome. Whereas for all DNA-seq libraries in the *fs(1)GS06* complementation group, there was a clear loss of read coverage at the insertion site, indicative of an insertion of an exogenous DNA sequence. Regardless, we were still unsure as to why we recovered this *Doc* insertion from our screen at such a high frequency.

We believe there are two possibilities for why we recovered this *Doc* insertion at *Grip91* at such a high frequency. The simplest explanation is that we were ‘floating’ this allele within our control stock at a relatively low frequency. We took measures to prevent this before the screen by isolating a single X-chromosome and then inbreeding the isogenized X with single, brother-sister matings, for six generations. If this insertion was indeed being floated, then it would have inserted after isolating a single X-chromosome and prior to, or during, the screening process. It is therefore possible that when we mated our ‘control’ *y*^*1*^ *cv*^*1*^ *v*^*1*^ *f*^*1*^ males to *flam*^−^ females, we eventually isolated chromosomes that previously contained the *Doc* insertion at *Grip91*. This would explain why we detected these reads within our control *y*^*1*^ *cv*^*1*^ *v*^*1*^ *f*^*1*^ DNA-seq libraries, especially if a few of the flies that were processed for library preparation contained this insertion.

A second possibility is that the *Grip91* first intron contains the remnants of a *Doc* insertion, hence why we detect reads that map to the *Doc* transposable element at this location in our control *y*^*1*^ *cv*^*1*^ *v*^*1*^ *f*^*1*^ flies. Based on the frequency of *Doc* insertions into *Grip91* recovered from our screen, this would therefore be a ‘hotspot’ for *Doc* insertions. We have evidence for *Doc* insertions in other alleles from our screen so we are confident that *Doc* transposable elements are being mobilized in *flam*^−^ females. We also detected *Doc* transposable element reads within the *Grip91* first intron in *fs(1)GS04, fs(1)GS05, fs(1)GS10*, and *fs(1)GS12*, but not other *fs(1)* alleles. This provides further evidence for the theory that there are indeed the remnants of a *Doc* transposable element at this location and that there may be a preference for *Doc* insertions in this region. Regardless, this complementation group highlights one of the main drawbacks of this screening method, which is the high frequency of unintended transposable elements insertions other than *mdg4*.

*fs(1)GS16* contained a novel *mdg4* insertion within the intron of *CG42699*, however, this is outside of the interval responsible for sterility.

#### fs(1)GS08

*fs(1)GS08* sterility mapped to the *cv*-*v* interval and we detected a novel *mdg4* insertion 233 NTs upstream of the gene *fs(1)M3. Df(1)BSC654* (containing *fs(1)M3*), *fs(1)M3*^*1901*^, and *fs(1)M3*^*K1*^ (Komitopoulou et al., 1983; Mohler & Carroll, 1984) all failed to complement *fs(1)GS08* sterility. Therefore, the upstream *mdg4* insertion was disrupting the activity of *fs(1)M3. fs(1)M3* is a known female sterile with requirement in the germline (Ambrosio et al., 1989; Degelmann et al., 1990; Perrimon & Gans, 1983). *fs(1)M3*, along with *fs(1)N* and *clos*, encode required components of the eggshell (Mineo et al., 2017; Ventura et al., 2010). Mutants for any of these genes result in perturbed eggshell formation and maternal effect lethality. The *mdg4* insertion upstream of *fs(1)GS08* is within an OVO ChIP peak, GSC, and St. 5 NC ATAC peak and roughly 85 NTs away from an OVO DNA binding motif.

*fs(1)GS08* was particularly interesting due to the upstream location of the *mdg4* insertion. Our previous work on OVO showed that *fs(1)M3* was directly bound by OVO and responded transcriptionally to OVO *in trans*, indicating it is likely regulated by OVO in the female germline (Benner et al., 2024). OVO binding is enriched at the TSS of *fs(1)M3* and there is a second enriched OVO bound peak upstream of the TSS. The *mdg4* insertion in *fs(1)GS08* is in between these two strong OVO peaks and due to *mdg4*’s known insulator activity, we believe that this insertion is evidence for a specific disruption of the upstream OVO bound region (Figure 1C). Whether this region is an active enhancer and is required for *fs(1)M3* expression in the female germline would be of interest for future work.

We also detected a novel *Doc* insertion along this chromosome to the right of *v* within the 3’UTR of *disco*. This insertion is outside the mapped sterility interval and therefore is not contributing to the sterility phenotype.

#### fs(1)GS10

*fs(1)GS10* sterility mapped to the *cv*-*v* interval. We did not detect any novel transposable elements within this region or along the rest of the chromosome. We refined the mapping interval to be to the right of *singed*, however, we do not have any further evidence for the precise disruption leading to female sterility. It is possible that this is an example of an unstable transposable element insertion, and within the time of isolating the allele and collecting samples for DNA-seq, the insertion had re-mobilized and been lost. A random spontaneous mutation other than a transposable element along this chromosome is also possible.

#### fs(1)GS11

*fs(1)GS11* sterility mapped to the *y*-*cv* interval and we found a novel *mdg4* insertion within the first intron of *VhaAC39-1*, commonly known as *chocolate*. A deficiency of this region, *Df(1)BSC886*, failed to complement sterility of *fs(1)GS11* females. *fs(1)GS11* complemented lethality of two recessive lethal alleles of *VhaAC39-1, VhaAC39-1*^*FY38*^ and *VhaAC39-1*^*FZ29*^ (Yan et al., 2009), but these transheterozygous females were sterile. A duplication of this region, that fully duplicates the genes *lncRNA:roX1, yin, CG2930, VhaAC39-1, CG15239*, and *CG42541*, rescued fertility in homozygous *fs(1)GS11* flies. Therefore, this is good evidence that this *mdg4* insertion is disrupting *VhaAC39-1* function resulting in female sterility. Mutations in *VhaAC39-1* have previously been shown to be required for *Notch* signaling in follicle cells of the ovary (Yan et al., 2009), however, there has not been a previously described role for *VhaAC39-1* in the germline. The *mdg4* insertion at this location is within a GSC and St. 5 NC ATAC peak, but not within an OVO ChIP peak. However, OVO does show significant binding to the promoter of *VhaAC39-1* with the closest DNA binding motif roughly 575 NTs away from the *mdg4* insertion site.

#### fs(1)GS12

*fs(1)GS12* sterility mapped to the *v*-*f* interval and we identified a novel *mdg4* insertion within the first intron of the gene *eIF5*. Deficiency *Df(1)BSC714*, containing *eIF5*, failed to complement sterility of *fs(1)GS12* and therefore this *mdg4* insertion was likely the cause of sterility in *fs(1)GS12* females. Knockdown of *eIF5* in the germline has previously been shown to result in germ cell death and sterility in adult females (Liu & Lasko, 2015). This *mdg4* insertion overlapped a GSC ATAC peak only and was within 340 NTs of an OVO DNA binding motif. However, there was no significant OVO binding within this region according to OVO ChIP-seq data.

#### fs(1)GS13

Meiotic mapping of *fs(1)GS13* was somewhat inconclusive. We found that *fs(1)GS13* recombinants *cv*^*1*^ *v*^*1*^ *f*^*1*^ and *v*^*1*^ *f*^*1*^ had both female fertile and sterile scoring recombinants, while all *f*^*1*^ recombinants were sterile. This result might indicate that the locus responsible for sterility is to the right of *f* and that the *cv*^*1*^ *v*^*1*^ *f*^*1*^ and *v*^*1*^ *f*^*1*^ fertile chromosomes were the result of double crossovers, with the second recombination event taking place in-between *f* and the female sterile locus. Consistent with this, we found a novel *mdg4* insertion to the right of *f* that was 120 NTs upstream of the pacman (*pcm*) TSS. *Df(1)Exel7468*, containing *pcm*, failed to complement sterility of *fs(1)GS13* females. Alleles *pcm*^*14*^ complemented *fs(1)GS13* while 50% (5/10) of *pcm*^*15*^/*fs(1)GS13* females were sterile. Of the 5 fertile *pcm*^*15*^/*fs(1)GS13* females, we found that after 5 days of egg laying, no more than 10 larvae had hatched in each individual vial (2, 2, 2, 7, and 8 hatched, respectively). This indicates that there was a non-penetrant maternal-effect phenotype in *pcm*^*15*^/*fs(1)GS13* females. *pcm* has previously been shown to be required for full female fecundity (Lin et al., 2008), so the weak fecundity phenotype of *pcm*^*15*^/*fs(1)GS13* females is not inconsistent with the phenotype previously described for *pcm. pcm*^*14*^ and *pcm*^*15*^ are both described as amorphic alleles, with different molecular lesions at the locus (Jones et al., 2016; Waldron et al., 2015). *pcm*^*14*^ contains a 3’ deletion of the open reading frame and *pcm*^*15*^ is an imprecise *P-element* excision containing residual *P-element* sequence within the 5’UTR. Since *pcm*^*15*^ and *fs(1)GS13* both contain exogenous DNA insertions upstream of the open reading frame, this could explain the low fecundity/sterility phenotype of respective transheterozygous females. Why *pcm*^*14*^ is able to complement *fs(1)GS13* is puzzling, however, a simple transvection model (Fukaya & Levine, 2017) of these two alleles could possibly explain the complementation result.

This insertion is within an OVO ChIP, GSC, and St. 5 NC ATAC peak while the closest OVO DNA binding motif is roughly 3,750 NTs away.

There was also evidence for a *Doc* insertion in the third intron of the gene *spri* (*spri-RN*) along this chromosome. This insertion is to the left of *v* and therefore not within the sterility interval based on the meiotic mapping results.

#### *mdg4* Insertions and OVO DNA Binding

Our original motivation for carrying out this screen was to recover *mdg4* insertions that caused female sterility based on the evidence that *mdg4* transposable elements insertions are in close proximity to OVO DNA binding motifs, and presumably, is where OVO is binding and regulating target gene expression. For the 13 novel *mdg4* insertions we detected, 5 were within a region that showed significant OVO DNA binding. Previously, *mdg4* insertions into the *ovo* promoter have been shown to cluster within a 200 NT window (Dej et al., 1998; Labrador & Corces, 2001), however, we found that only 3/13 *mdg4* insertions were within 200 NTs of an OVO DNA binding motif. Labrador et al., 2008 showed that a synthetic DNA construct containing OVO DNA binding motifs had a significantly larger window for *mdg4* insertions (up to roughly 900 NTs) outside of the OVO DNA binding motif array and the authors suggested that mechanisms in addition to OVO binding (e.g. transcription, nucleosome position, open chromatin, etc.) are likely driving *mdg4* insertion bias. A majority of the *mdg4* insertions (10/13) recovered from our screen are > 200 NTs away from an OVO DNA binding motif and therefore supports the authors conclusions that other mechanisms are driving *mdg4* insertion bias. The *ovo* promoter likely contains a unique repertoire of these features/elements that drive *mdg4* insertions into such a narrow window.

Since only 5/13 *mdg4* insertions were within a region bound by OVO, it is possible that the relationship between OVO binding and *mdg4* insertion might not be as strong as previously reported. For example, Labrador et al. 2001 provided evidence that the quantity of *mdg4* insertions into the *ovo* promoter are much higher in an *ovo*^*D1*^ background than when compared to an *ovo*^*+*^ background. This result might indicate that the increased quantity of OVO-A-like protein expressed from *ovo*^*D1*^ might be driving the OVO-*mdg4* relationship more specifically, whereas the predominant OVO-B protein in *ovo*^*+*^ females has a weaker relationship with *mdg4* insertional bias. To test this, we randomly shuffled the locations of significant OVO ChIP peaks 1000 times and intersected the shuffled peak locations with all the *mdg4* insertions recovered from our screen. Since 5 insertions were found to be within an OVO ChIP peak, we could then test if there was an enrichment for *mdg4* to transpose into OVO bound regions by counting the number of times the randomly shuffled sequences intersected 5 or more *mdg4* insertions. We found that out of the 1000 shuffled OVO ChIP peak locations, zero had 5 or greater *mdg4* intersections, and therefore, this result suggests that there is indeed a significant enrichment for *mdg4* to insert into OVO bound regions (p < 0.001) and the propensity of *mdg4* to insert within an OVO bound region is unlikely a result of the background level of OVO binding in the genome.

With all the combined evidence from our pilot screen, we ultimately decided against expanding this screen for direct OVO target genes in its current format. The primary reason is that we were only interested in mutagenesis via *mdg4* transposition because of its previously described insertional bias near OVO DNA binding motifs. We do have evidence for a strong OVO-*mdg4* relationship in an *ovo*^*+*^ background, however, *mdg4* insertions only accounted for 8/17 (47%) female sterile alleles recovered. We would therefore have to double our efforts to recover an appreciable amount of female sterile alleles due to *mdg4* in this current format. Also, in seven of our alleles, there were two or more novel transposable element insertions along a single chromosome. Another unwanted consequence of this method.

We do believe that this method potentially has value, outside of the reasons stated above, with a few alterations. Reading frames have been transgenically incorporated into *mdg4* sequences allowing for the detection of *mdg4* activity in permissive cellular conditions (Chang et al., 2019). This method could be utilized for screening purposes by incorporating a reporter into a full-length *mdg4* element. In our case, this full length *mdg4* element including the reporter could then be mobilized in *flam*^−^ females and one could isolate only flies where the *mdg4* reporter has now transposed to the chromosome of interest. This would speed up the screening process significantly since you would only test chromosomes for sterility that have a novel *mdg4* insertion identified via the presence of the reporter. One would also likely recover fewer chromosomes of interest due to insertions of non-*mdg4* transposable elements, but the possibility still clearly exists, so meiotic mapping would still need to be completed to confirm. There are also alternative methods to isolating *mdg4* induced mutations that might possibly reduce the background of non-*mdg4* mutation rates. For example, it has been shown that wild type larvae cultured in media supplemented with *flam*^−^ pupae homogenate, or isolated viral extracts, can induce *mdg4* transposition in the next generation (Dej et al., 1998; Kim et al., 1994; Song et al., 1994). Presumably, active *mdg4* retroviral particles are maternally deposited by these otherwise wild type flies. However, it is unknown if other transposable elements are also maternally deposited with this method.

Regardless, this approach still has utility based on the insulating properties of *mdg4* elements and the ability to disrupt enhancer-promoter communication. 3/17 of our alleles were intergenic, indicating that *mdg4* was likely disrupting an enhancer’s ability to influence gene expression of its target gene. With a genetically modern and rational design of a self-reporting *mdg4* retrotransposon, we believe this screening method could be of great value for scientists especially interested in dissecting cis-regulatory logic and transcription, and more generally to those interested in mobile genetic elements.

## Data Availability Statement

DNA-seq data generated in this study are available at SRA:PRJNA1422399. Female germline ATAC-seq datasets analyzed in this study are available at SRA:PRJNA606593. OVO ChIP-seq datasets can be found at SRA:PRJNA1041436.

**Figure S1: Cross scheme and reversion rates of *ovo*^*D1*^.**

**Table S1: Flybase ART Table.**

**Table S2: Meiotic mapping and complementation test results.** The interval that female sterility mapped to based on meiotic mapping using the phenotypic markers *y*^*1*^, *cv*^*1*^, *v*^*1*^, and *f*^*1*^. Black boxes indicate that the alleles were not complementation tested, green boxes denote alleles that complemented each other, red boxes denote alleles that failed to complement each other.

**Table S3: Characteristics of all non-*mdg4* dependent recessive female sterile alleles.**

## Acknowledgements

We thank Norbert Perrimon for supplying reagents used in this study. Stocks obtained from the Bloomington Drosophila Stock Center (NIH P40OD018537) were used in this study. Sequencing was performed by the DNA Sequencing and Genomics Core at the National Heart, Lung, and Blood Institute (NHLBI) and the National Institute of Diabetes and Digestive and Kidney Diseases (NIDDK) Genomics Core. Genetic and genomic information was obtained from the UCSC Genome browser and FlyBase (U41 HG-000739). This work utilized the computational resources of the NIH High-Performance Computing Biowulf cluster (http://hpc.nih.gov).

